# Extracellular vesicles released from cortical neurons influence spontaneous activity of recipient neurons

**DOI:** 10.1101/2025.04.09.647922

**Authors:** Franco Luis Lombino, Mohsin Shafiq, Andreu Matamoros-Angles, Jürgen R. Schwarz, Kira V. Gromova, Daniele Stajano, Bente Siebels, Leonie Bergmann, Tim Magnus, Michaela Schweizer, Franz Lennard Ricklefs, Hartmut Schlüter, Andrew F. Hill, Matthias Kneussel, Markus Glatzel

## Abstract

Extracellular vesicles (EVs) are membranous structures that cells release into the extracellular space. EVs carry various molecules such as proteins, lipids, and nucleic acids, and serve as specialized transporters to influence other cells.

In the central nervous system, EVs have been linked to many important processes, including intercellular communication, but molecular details of their physiological functions are not fully understood. Our study aimed to investigate how EVs are released by neuronal cells, and how they affect the neuronal activity of other recipient neurons. We show that mature primary cortical neurons release EVs from both their soma and dendrites. EVs released from neurons closely resemble non-neuronal EVs regarding size and marker proteins and proteomic analyses showed that neuronally released EVs contain proteins typically acting in pre- and post-synaptic compartments. Interestingly, our analysis revealed that EVs alter spontaneous activity in target neurons by increasing the amplitude of postsynaptic potentials.

In summary, our findings elaborates on the role of EVs in synaptic activity modulation in neurons mediated by glutamate receptors.

## Introduction

Extracellular vesicles (EVs) are released into the extracellular space by every cell type^1^. They are a population of distinct membranous units and carriers of different types of biological material. Indeed, it is known that nucleic acids, proteins, and lipids can be transferred to cells using EVs as shuttles^2^. EVs, mainly consisting of exosomes and microvesicles, are thought to have biologically relevant functions in cell-to-cell communication, and their therapeutic potential is of great interest^3^. Even if the molecular composition of specific EV subtypes is not fully uncovered, their molecular identity is thought to depend on the physiological state of the donor cells and the existence of specific tetraspanins in the EV membrane, which are also linked to specific functions of EVs^4,5^.

In the brain, EVs are secreted by neurons, astrocytes, oligodendrocytes and microglia^6^. Interestingly, EVs from different sources appear to have different functions in target cells^7^. In the central nervous system, EVs play key roles in brain development^8^ and are directly involved in the spreading of pathogenic molecules in neurodegenerative diseases^9^.

Chemical synapses are central players in neuronal communication and recent data indicate that EVs play roles in some aspects of synapse physiology. For instance, EVs derived from primary astrocytes impact neuronal synapse development^10^ and the transfer of synaptic proteins such as synaptobrevin between neurons occurs via EVs^11^. Furthermore, AMPA (α-amino-3-hydroxy-5-methyl-4-isoxazolepropionic acid) receptors are present in EVs, with neuronal activity regulating their secretion^12^. Despite this evidence, the functional roles that neuronal EVs exert in neighboring recipient neurons are not fully uncovered. To better understand whether and how EVs secreted from donor neurons modulate synaptic activity in recipient neurons we isolated and characterized cortical neuronal culture-derived EVs, assessed their protein composition by mass spectrometry and performed electrophysiological experiments in cultured hippocampal recipient neurons.

Here, we demonstrate that neurons release EVs not only from their soma but also from dendritic compartments. Moreover, we describe that the neuronal EVs enhance the amplitude of spontaneous post-synaptic potentials in recipient neurons. Furthermore, our electrophysiological and proteomic analyses provide molecular cues on how EVs modulate synaptic activity highlighting the role of glutamatergic signaling in this process.

## Results

### Cortex-derived primary cultures display active synapses and release extracellular vesicles

To evaluate the role of EVs in synaptic modulation, we used cortex-derived primary neuronal cultures from mouse embryos. To verify the cellular identity in our primary culture model, we stained DIV14-15 cultures with antibodies against both neuronal-specific enolase (NSE) as a neuronal marker and glial fibrillary protein (GFAP) as a marker for astrocytes. Immunostaining confirmed that our model represents a mixed culture consisting of primary neurons and astrocytes (Fig. 1A and Supplementary Fig. S1). Next, we tested if neurons displayed structures typically observed in mature neurons and co-stained the microtubule-associated protein 2 (MAP2) and the F-actin filaments with rhodamine-phalloidin to detect dendrites and dendritic spines. Indeed, our neurons contain synaptic compartments (Fig. 1B) that can be clustered into filopodia, stubby and mushroom spines (Fig. 1C). To test if cortical neurons at DIV14 were able to form mature synapses, we co-stained against synaptophysin (pre-synaptic marker) and the post-synaptic density protein 95 (PSD-95, post-synaptic marker) (Fig. 1D). The co-localization of both markers indicated the presence of mature excitatory glutamatergic synapses (Fig. 1E). In summary, our cell culture model has features of mature cortical neurons in co-existence with glial cells, being a suitable *in vitro* model to study the role of EVs in neuronal activity. To assess if primary cortical neurons spontaneously release EVs, we utilized a previously characterized CD63-pHluorin construct to monitor the fusion of the multi vesicular bodies (MVB) to the plasma membrane and GFP fluorescence as an indication of EV release^13^. In combination with a volume marker to observe cellular morphology, we performed TIRF microscopy. We distinguished two types of GFP clusters (Fig. 1F and G): membrane-stable clusters (Fig. 1G asterisk) and dynamic clusters representing release events (Fig. 1G arrowhead). As expected from a protein of the tetraspanin family, a significant fraction of CD63 remains immobile in the cell membrane likely in tetraspanin-enriched microdomains. However, MVB fusion and, therefore, EV release occurred both in the cell soma and dendritic structures.

**Figure 1.**
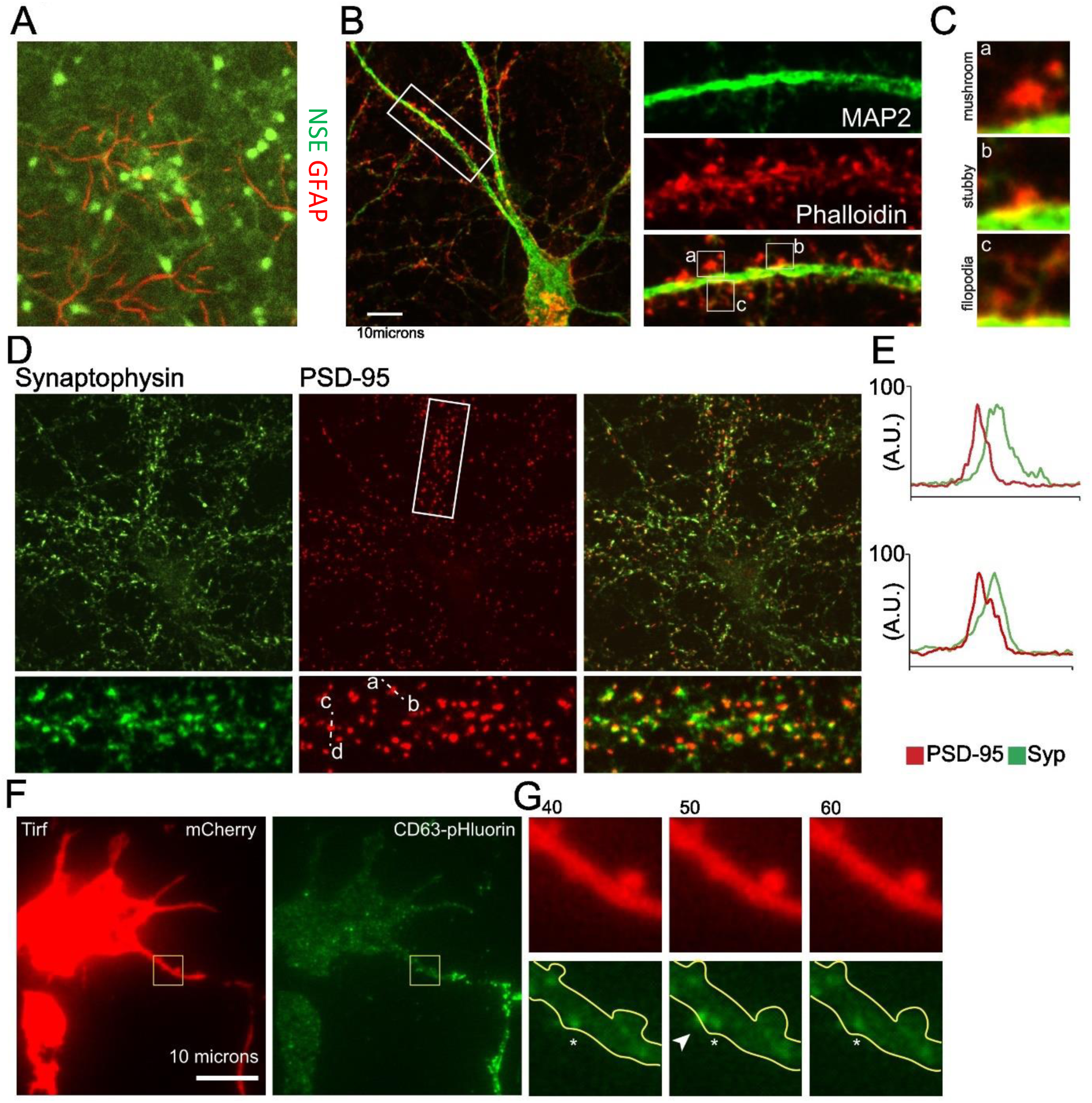
Cultured cortical neurons secrete extracellular vesicles. **A)** Immunofluorescence staining of primary neurons derived from mouse cortex. NSE (green) and GFAP (red) coexistence indicates the presence of both glial cells and primary neurons (n=3 independent experiments). **B)** Primary neurons co-stained with MAP2 and Phalloidin to observe postsynaptic spines in neuronal dendrites. Scale bar 10 microns (n=3 independent experiments). **C)** MAP2 and Phalloidin staining shows the presence of mushroom (a), stubby (b) and filopodia (c) spines in B. **D)** Co-staining of the presynaptic marker synaptophysin (Syp) and the postsynaptic marker PSD95 indicates the presence of active glutamatergic synapses. (n=3 independent experiments). **E)** Overlapping profile of the co-staining synaptophysin and PSD95 shown in D. A.U. is arbitrary units. **F)** Timelapse microscopy showing the overexpression of the volume marker mCherry and the pH-sensitive CD63-pHluorin plasmid as an indicator of MVB fusion to the plasma membrane and indirectly suggesting the release of exosomes from cell bodies and dendrites of cortical neurons. Scale bar 10 microns. (n=3 independent experiments). **G)** Magnification of region indicated in F. The star indicates a permanent patch of CD63-pHluorin in the plasma membrane, the arrow a brief exosomal release event.

From the same neuronal cultures, EVs were isolated and characterized following the MISEV2023 guidelines^14^. EVs were purified via differential centrifugation, and Nanoparticle Tracking Analysis (NTA) measurements showed an EV population with an average size of 137.6 ± 4.4 nm (Fig. 2A and B). Further characterization of EVs by western blotting with antibodies to GM130 (a Golgi apparatus marker, used as an EV purity marker) and EV markers CD81, Tsg101, and Flotillin-1 validated the purity of our EV isolation (Fig. 2C and Supplementary Fig. 2). Lastly, we performed transmission electron microscopy to further validate EVs’ morphology and shape, which showed the characteristic membranous EV cup shape (Fig. 2D). In summary, our EV preparations from mature cortex-derived primary neuronal cultures were highly enriched for intact EVs of 70-150 nm size in diameter which expressed membranous and intraluminal EV markers. This prompted us to further investigate the makeup and function of these EVs.

**Figure 2.**
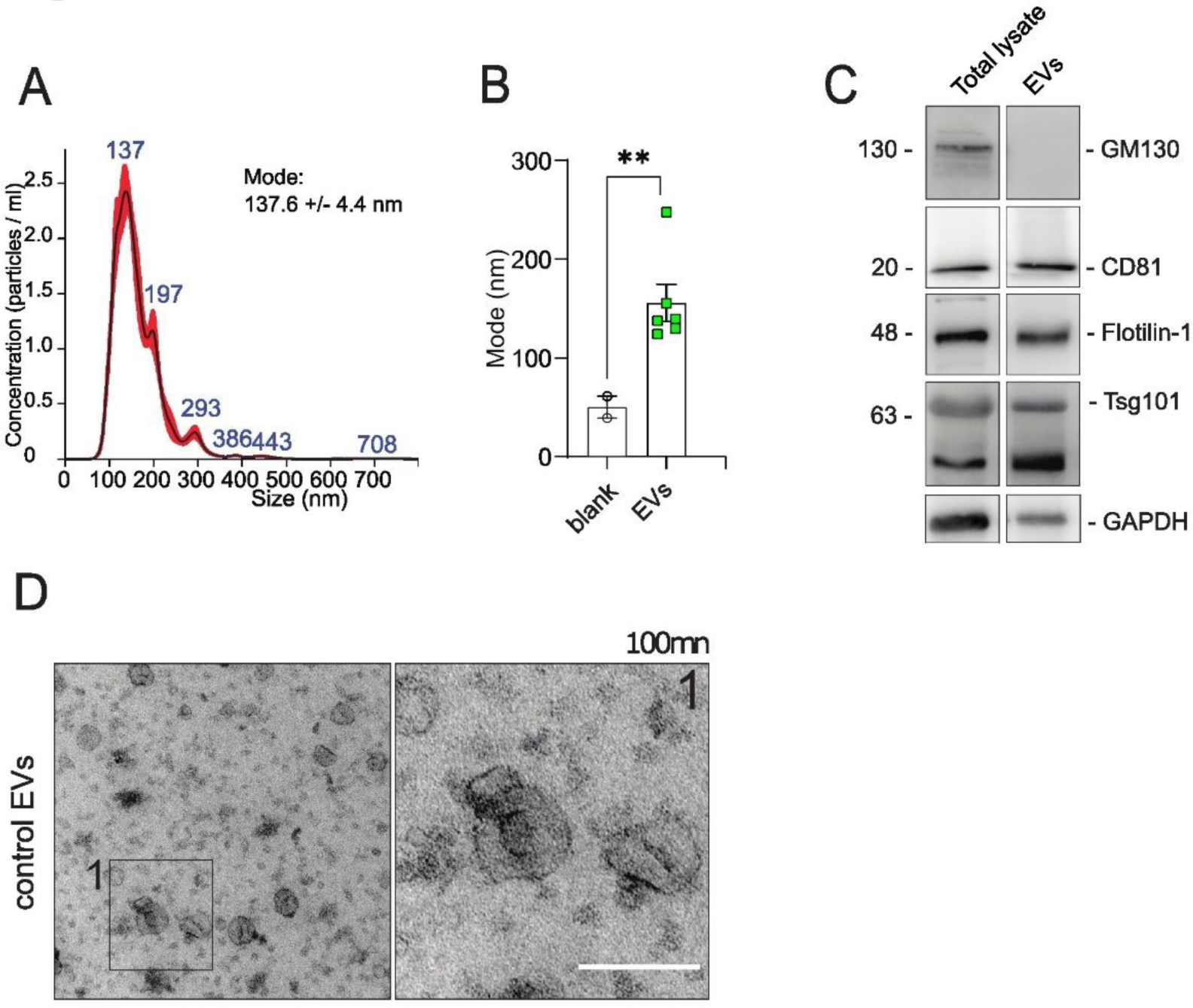
EV isolation from cultured cortical neurons. **A)** Nanoparticle tracking analysis representative example of extracellular vesicles isolated via differential centrifugation. **B)** Average mode size of EVs isolated from primary cultures reflects the expected particle size as compared to the control buffer. **C)** Characterization of isolated EVs by western blot shows the presence of markers typically present in EVs (CD81, Flotillin-1 and Tsg101) and the lack of GM130 (negative control). **D)** Electron microscopy of purified EVs showing the presence of particles of around 100nm in diameter as well as smaller vesicles. Scale bar, 100 nm

### The proteomic analysis of cortical culture-derived EVs reveals that they are carriers of neuromodulatory proteins

To assess whether neuronal EVs are enriched in proteins involved in synaptic function, we performed global proteomics of EVs and corresponding cell lysates (n = 6 in both) (Fig. 3). Except for 26 proteins found exclusively in EVs, we found 2,415 proteins overlapping between EVs and lysates, as shown in the Venn diagram that depicts all the detected proteins across samples (Fig. 3A). We also found a high degree of intragroup similarity between the lysate samples compared with the EVs, and vice versa, showing a higher correlation with the samples from the same origin (Fig. 3B and C). Moreover, the presence of several known EV markers shown in Figure 3D argues in favour of the enrichment and purity of our EVs preparations. The volcano plot (Fig. 3E) comparing the protein abundances in EVs preparations and lysates shows a variety of proteins significantly enriched in EVs, such as Adam10, Apoe and Ezr; (in orange) and those preferentially present in the donor cells’ lysates (in grey). Gene ontology (GO) analysis highlighting the biological process in which proteins preferentially found in EVs are involved, reveals roles primarily related to neuronal development, proliferation and differentiation (Fig. 3F, supplementary data: 2). Furthermore, we looked for the potential involvement of EV proteins in synaptic and intracellular signaling processes (Fig. 4) by performing GO (supplementary data: 3) and Reactome pathway enrichment analyses (supplementary data: 4). The GO enrichment analyses (Fig. 4A) argue in favor of EVs being involved in synaptic and trans-synaptic signaling, primarily driven by the presence of glutamate receptors (evident from the presence of glutamate receptors subunits GluA1, GluA2 and GluN2b in the data) and their interactors CamkIIa and Gsk3b (Fig. 4B and C). Interestingly, the presence of AMPA-receptors and NMDA receptors in our EVs could be confirmed by western blot of EV fractions derived from neuronal cultures (Fig. 5A). Moreover, using immunoelectron microscopy, AMPA receptor GluA2 could be mapped to the outer leaflet of the EV lipid bilayer (Fig. 5B). To assess the *in vivo* relevance of our data, we evaluated the presence of AMPA and NMDA receptors in brain-derived EVs (BDEVs) obtained from mouse brain^15^. The BDEVs were of high purity as shown by NTA and TEM (Supplementary Fig. S3). These analyses confirmed the presence of AMPA-receptor subunits (GluA1, GluA2) and NMDA receptor subunit Nr2b on EVs (Fig. 5C).

**Figure 3.**
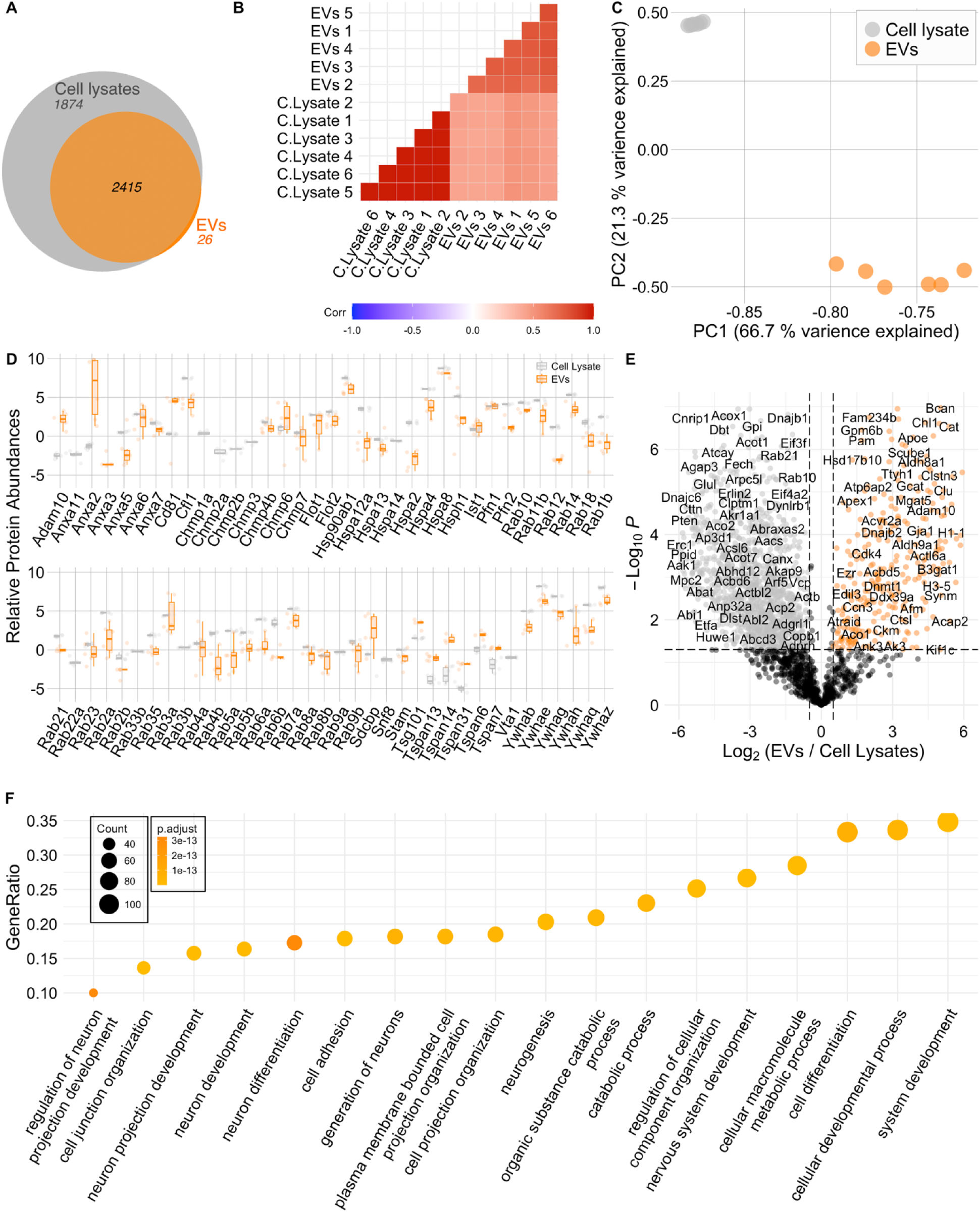
Proteomic composition of EVs derived from Primary neurons. A) Venn diagram depicts a major overlap in proteomic composition of EVs sourced from primary neurons to that of respective cellular lysates. The correlation plot (B) (with high Pearson’s correlation values) and Principle Component Analysis - PCA (C) underscore the internal consistency within the EV samples and lysates, highlighting the degree of similarity in their protein expression patterns. Calculation of correlation values and PCA were conducted for the proteins which were commonly expressed between the EVs and lysates. D) Relative abundances of various EV marker proteins in the EVs and Lysates are depicted graphically. The abundance of these markers confirms the purity of the isolated EV population, crucial for accurate interpretation of subsequent analyses. E) Volcano plot showing upregulated and downregulated proteins in EVs compared to Lysates depicting the proteins preferentially packed in the EVs (highlighted orange) with respect to the donor lysates. F) The dotplot showcases the top gene ontology terms (category: biological processes), that are linked to the proteins predominantly found within EVs.

**Figure 4:**
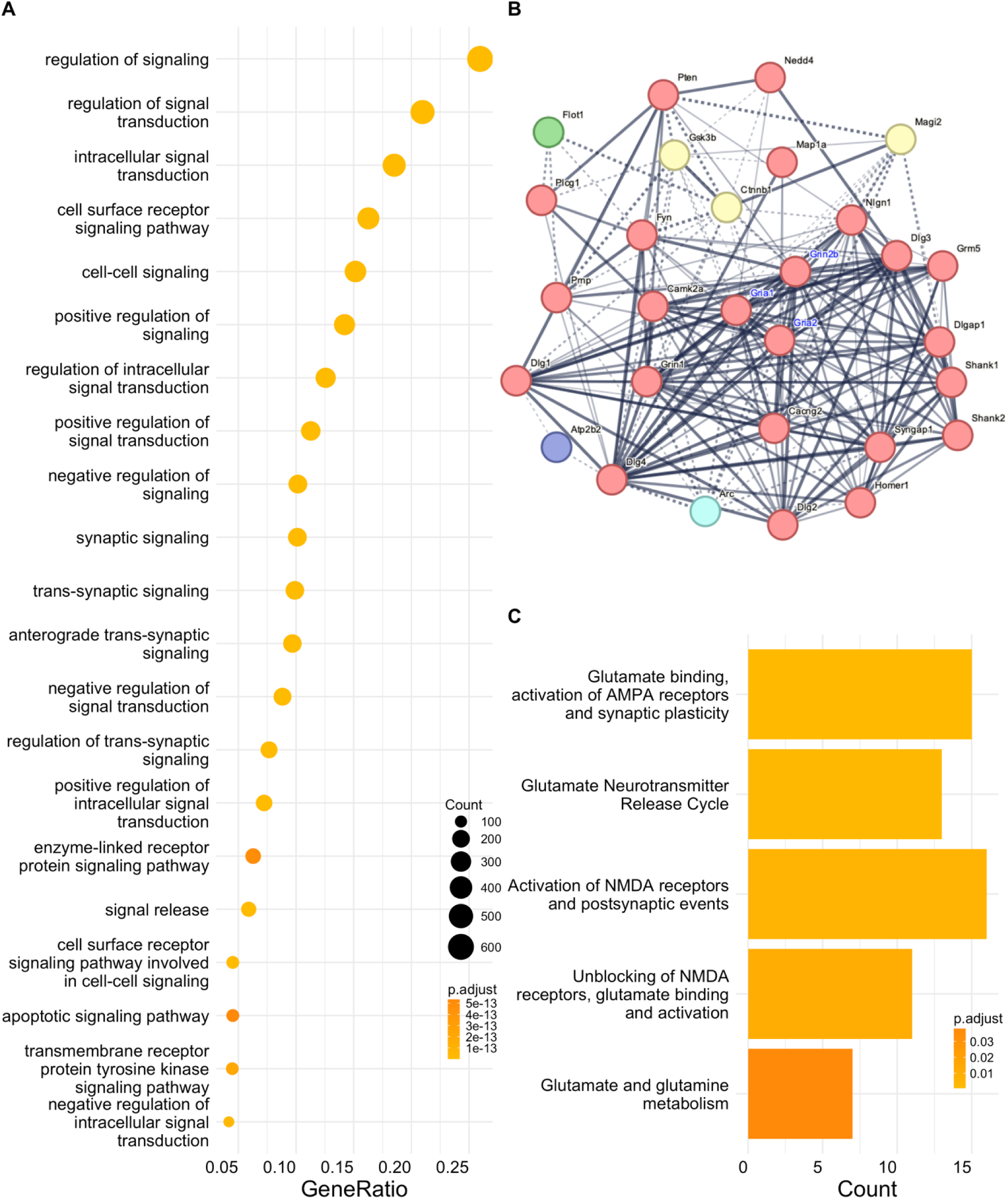
Certain proteins in primary culture derived EVs were associated to cell-cell signaling, trans-synaptic signaling, and NMDA receptor trafficking and activity. A) The dot plot presented here illustrates diverse signaling (Synaptic and receptor protein kinase) pathways identified through the GO enrichment analysis. List of the proteins is presented in supplementary data: 3. B) Network of the EV proteins (from the proteomics data) related to Glutamate receptor binding pathway is shown here. Network was assorted together with String-db application in cytoscape. C) Reactome pathway enrichement for the proteins expressed in the EVs shows their active involvement in the AMPA receptor and NMDA receptor trafficking depicted here by dotplot (supplementary data: 4).

**Figure 5.**
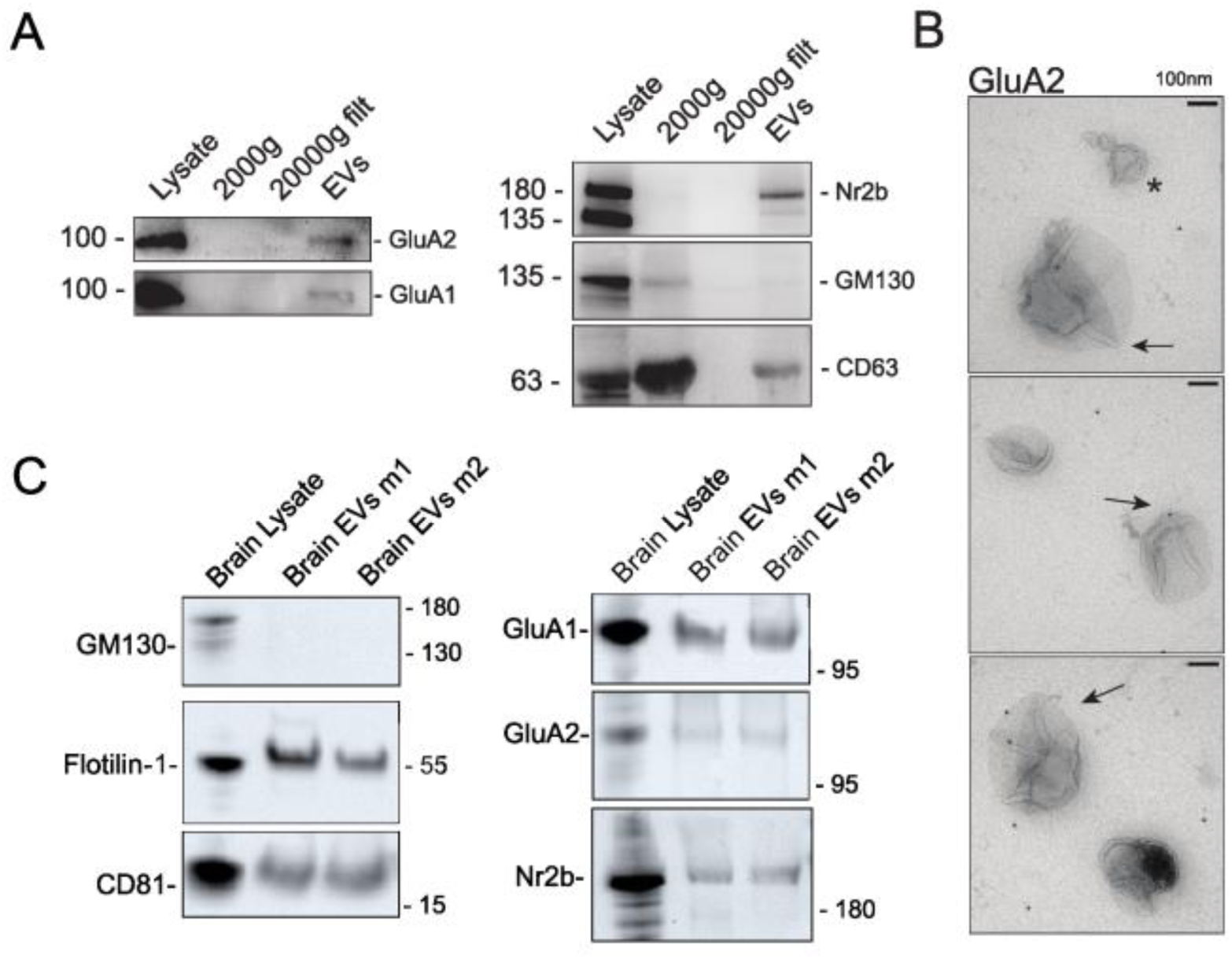
EVs isolated from primary cultures contain AMPA receptor subunits. **A)** Glutamate receptor subunits are present in EVs isolated from primary cultures, in combination with the EV-marker CD63. **B)** Immunoelectron microscopy of EVs isolated from cultured neurons support the presence of the glutamate receptor subunit GluA2. **C)** EVs isolated from mouse brains and validated qualitatively by the presence of the EV markers Flotillin-1 and CD81, contain both AMPA receptor subunits and Nr2b subunit of NMDA receptors.

### EVs derived from cortical cultures modulate the synaptic activity of recipient hippocampal cultures

Aiming to understand the role of neuronally released EVs in synaptic activity, we applied EVs derived from primary cortical cultures to DIV14 hippocampal neuron cultures. After overnight incubation (Fig. 6A), we performed current-clamp recording and evaluated the frequency of spontaneously firing action potentials and the amplitude of postsynaptic potentials. As shown in Fig. 6B-D, EVs did not alter the frequency of synaptic potentials when compared to the no-EV control (Fig. 6B and C), but led to significantly increased amplitudes in post-synaptic potentials (PSP) (Fig. 6B and D). Additionally, the application of DNQX (a potent and selective AMPA/kainate-receptor antagonist) to the recipient neurons markedly reduced neuronal activity, and subsequent washing out of the DNQX led to a reactivation of the synaptic activity (Fig 5 E, F). Together, these results indicate that EVs are synaptic modulators and influence post-synaptic potentials in recipient cells.

**Figure 6.**
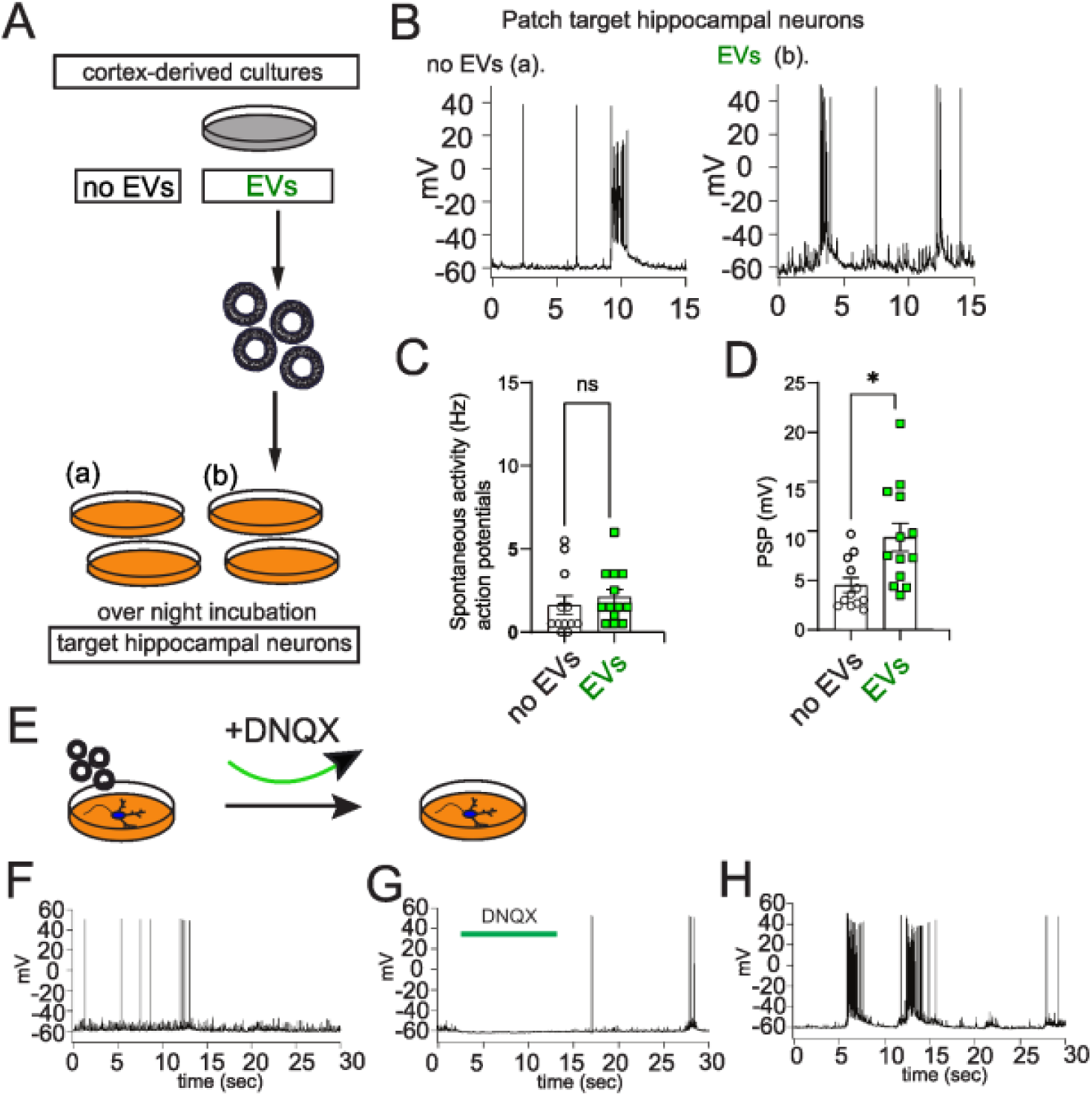
Extracellular vesicles modulate the synaptic activity of target cells. **A)** Strategy and conditions of EVs administration protocol for electrophysiological recordings. Briefly, EVs from cultured cortical neurons were administered to hippocampal primary cultures overnight. Patch clamp recordings were performed the day after. **B)** Example traces of patch clamp recordings of hippocampal cultures. Left panel “a” shows a recording from a neuron of hippocampal cultures which received only vehicle solution (PBS/inhibitors). Panel “b” in the right shows a recording of a neuron of hippocampal cultures which received EVs from non-stimulated cortical cultures. **C)** Quantification of frequency of action potentials show no significant difference between conditions. (n=3 independent experiments). **D)** Amplitude of events is significantly affected by the presence of EVs, irrespective of glycine in the donor cortical culture (n=3 independent experiments). E-H) Shows that application of DNQX abolishes postsynaptic potentials and its subsequent washing leads to a re-activation of synaptic activity, implicating that primarily excitatory currents were recorded.

In summary, our data strongly argue in favour of a prominent contribution of EVs in modulating synaptic activity in recipient cells, and based on our mass spectrometry data, we propose involvement of glutamate receptors in this process.

## Discussion

EVs have been related to numerous cellular processes in the central nervous system^16^. Likely because of the diversity of cells that secrete functionally different subtypes of EVs^17^, their impact on target cells can be highly heterogeneous. For instance, lipids transported in EVs are known to alter target cell membrane characteristics^18^. Moreover, proteins contained in EVs can regulate EVs biogenesis, secretion, or uptake by target cells^17,19^. Here we show that secretion of EVs occurs at multiple sites within neurons, including dendrites and soma. Although in dividing cells, this may have minor implications, in large and highly polarized neurons, characterized by a soma, a complex dendritic tree and an axon, each one with highly diverse molecular and electrical properties, secretion from different compartments may affect the molecular make up and function of the secreted EVs^20,21^. In this respect, it was shown that MVBs, the source of the exosomes, are widely distributed in both soma and dendrites of neurons *in vivo* and *in vitro*^22^.

In this study, we revealed that EVs impact the amplitude of post-synaptic potentials (PSP) or target cells, and the blockage of neuronal activity via DNQX, suggests that the major component transferred by cortical EVs was excitatory, from the glutamatergic system. Similarly, it was recently shown that vesicles released by glioma cells impact synchrony in neurons^23^. In this study, the authors showed that in addition to increasing Arp3 protein levels, exosomes derived from glioma cells increased the number of excitatory synapses in recipient neurons. In another study, the authors characterized the roles of exosomes and the larger microvesicles in modulating the spontaneous activity of cortical neurons^24^, and although a complete understanding of the different roles of these two populations in target cells was not achieved, the study adds additional evidence to the role of EVs in modulating brain function. How do EVs lead to an increase in PSP? Uptake of BDNF and TrkB receptor-containing EVs by clathrin-dependent endocytosis by neurons has been shown to modulate synapse density through TrkB receptor activation^25^. Thus, it is conceivable that the equilibrium of glutamate receptors responsible for fine-tuning post-synaptic potentials in neurons is altered by the uptake and membrane integration of glutamate receptors contained in EVs^26^. Along the same line, recent work showed that neuron-derived EVs increase excitatory synapse formation also in hippocampal neurons due to the delivery of certain miRNAs by the EVs, which could also explain our results^27^.

Aiming to uncover molecular contributors mediating synaptic changes in target hippocampal neurons, we performed mass-spectrometry analysis of EVs and lysates from donor cortical neurons. Interestingly, we identified a variety of proteins involved in neuronal development and pre- and post-synaptic regulation in EVs. These include proteins responsible for modulating post-synaptic potentials. In parallel, our results show that EVs from cortical neurons contain AMPA and NMDA receptors. Others have shown that AMPA receptors are frequent cargoes of EVs^28,29^, and we confirmed that both are present via western blot and immunoelectron microscopy. In parallel, we supported our results by showing the presence of both AMPA and NMDA receptor subunits in EVs obtained from mouse brain tissue. In recent work, the proteomic analysis of brain derived-EVs from mouse and human tissue revealed a prominent expression of proteins involved in postsynaptic density and specialization and similar synaptic functions^30^. Data on the presence of NMDA receptor subunits in EVs is scarce. One study by Fauré and others^28^ observed the presence of AMPA receptor subunits, but not GluN1 subunit of NMDA receptors in cortical cultures derived EVs. Others have found GluN1 subunit and other glutamate receptors in EVs in exosomes derived from human plasma^31,32^. GluN2b is likely in distinct vesicles that are a part of those containing AMPA receptor subunits. Since NMDA receptors are implicated in neuronal maturation, our findings highlight the putative role of EVs in modulating neuronal development and maturation^33^.

While our study is subject to limitations inherent to *in vitro* models, including the potential confounding effects of astrocyte-derived EVs^34^, it advances our understanding of EVs as modulators of neural communication and brain function. Specifically, our findings demonstrate that EVs modulate specific components of synaptic activity and contribute to the maintenance of target neuron synaptic properties. Furthermore, we provide novel insights into the potential role of EVs in the cross-regional regulation of synaptic activity, suggesting that EVs derived from cortical primary neurons can influence synaptic function in hippocampal primary neurons.

## Methods

### Mice

Animals were housed under controlled environmental conditions according to all ethical regulations in agreement with the European Community Council Directive (2010/63/EU), reviewed and approved by the ethics committee of Hamburg (Behörde für Justiz und Verbraucherschutz, Fachbereich Lebensmittelsicherheit und Veterinärwesen) and the animal care committee of the University Medical Center Hamburg-Eppendorf.

### Cortex-derived neuronal cell cultures

Mouse embryos were extracted from the uterine horns of pregnant female mice at E.16 and brains were harvested. Subsequently, cortices were isolated and incubated for 5 minutes at 37°C in the presence of 0.005% Trypsin to promote dissociation. Later, trypsin was blocked with FBS/HBSS, and all substituted by HBSS. Finally, further dissociation was achieved by a few passages through a glass pipette. For EV isolation, 0.7 cortices were plated on each 10 cm plate coated with poly-L-lysine the day before. In total, 6 to 10 plates were prepared for each isolation. For immunocytochemistry, cells were seeded on 10 mm glass coverslips previously coated with poly-L-lysine.

### Immunocytochemistry

Cortex-derived primary cultures were fixed after DIV12 using 4% formaldehyde/PBS for 3-5 minutes. Later, cells were washed 3 times with PBS, permeabilized for 3-5 minutes with Triton X-100/PBS and washed further with PBS for 3 additional times. Subsequently, 1% BSA/PBS was used for blocking 30-60 minutes at room temperature. Primary antibodies were incubated in blocking solution for 1 hour and washed 3 times. Finally, the secondary antibody was incubated in a blocking solution and, after washing thoroughly, coverslips were mounted with Aqua Poly-Mount. Images were obtained with a spinning disk microscope and processed using Fiji.

### Time lapse TIRF microscopy

Cortical cultures were seeded on glass coverslips as above and maintained in culture until young stage. Cells were transfected at DIV 6-7 with Calcium Phosphate method using CD63-pHluorin construct from addgene (#130901) and mCherry as a volume marker. One day later, after replacing the media with fresh Hepes buffer, cells were imaged at the spinning disk microscope in the presence of 5% CO_2_ and controlled temperature. Images were acquired applying Tirf for 200 frames with an interval of 2 seconds, then analyzed on Fiji.

### Extracellular vesicle isolation from primary cultures

EVs were isolated by a protocol developed by others^35^. In detail, cortex-derived primary cultures were kept in culture for 12-14 days in vitro in 10 cm cell culture dishes, previously coated with poly-L-Lysine. On the experimental day, media was collected and kept on ice in the presence of protease inhibitors. Cells were collected after scratching in ice-cold Triton/PBS supplemented with protease and phosphatase inhibitors. Cell lysates were centrifugated for 10 minutes at 4°C at 1,000xg. Supernatant was conserved for further analysis. Media was centrifugated 2,000*x*g at 4°C for 10 minutes. Later, the supernatant was collected and centrifugated 20,000*x*g for 30 minutes at 4°C. Cell debris were removed by filtration with a 200 nm filter and finally, flow through was centrifugated for 90 minutes at 4°C in SW40-Ti swinging rotor. Supernatant was discarded. EVs were at the bottom of the tube and resuspended in filtered PBS containing protease inhibitors.

For *electrophysiological experiments,* the control condition was represented by primary hippocampal cultures administered with PBS containing protease inhibitors, in the tested condition, primary hippocampal cultures received EVs from primary cortical cultures resuspended in PBS plus protease inhibitors. All conditions received the same volume of material. Incubation was performed overnight in 300-400 microliters of the primary hippocampal cultures pre-conditioned media.

### Isolation of Extracellular vesicles from mouse brain

EVs were isolated from frozen brain tissues obtained from BL6 mice (n = 4; 35-50 weeks old) as previously described^15^. In brief, the tissue was mechanically minced and digested with DNAse I (40 U/ml) and Collagenase D (2 mg/ml, Roche) for 30 min at 37°C. To remove the cell and tissue debris, the samples were centrifuged at 300*x*g and 2,000*x*g. Then, the supernatant was centrifuged at 16,000*x*g to pellet the large EVs (lEVs), and later at 118,000*x*g, to pellet the small EVs (sEVs). The lEVs and the sEVs were mixed and centrifuged again in a three-layer Optiprep cushion (Stemcell) (45%, 30% and 10%) at 185,000*x*g. The EVs were obtained from the fraction between the 30% and 10% layers.

### Extracellular vesicle characterization using Nanoparticle tracking analysis (NTA) and western blotting

*For NTA:* EV pellets obtained as previously described were diluted 1:1,000 in sterile and filtered water for NTA measurements.

*For western blot:* EVs from different conditions were normalized based on Nanoquant measurements, diluted with Laemmli buffer containing beta-mercapto ethanol for all antigens, except for tetraspanins where urea-containing Laemmli buffer was used. Finally, samples were either boiled or warmed up to 65°C respectively, and loaded onto SDS-page.

### Electron microscopy and immunogold labeling

For pelleting, EVs from cortical cell cultures were ultracentrifugated at 100,000*x*g for 70 minutes. The procedure was performed as in (A. Matamoros-Angles et al, 2024^30^). Shortly, the EVs were resuspended in 2% paraformaldehyde (PFA) for fixation. A fraction of this solution (5 µl) was adsorbed for 20 minutes to glow discharged carbon-coated Formvar grids (EMS, Germany). After a brief washing step with PBS, samples were subjected to post fixation with 1% glutaraldehyde in PBS and incubated on ice-cold methylcellulose-uranyl acetate solution for 30 minutes. Grids were then looped out and air-dried before analysis at the electron microscope.

For immunogold labeling, EVs were absorbed on grids as described above. Then, they were rinsed in PBS and quenched in 20 mM glycine in PBS. 1% bovine serum albumin (BSA) was used as blocker for 10 minutes^36^. Grids with EVs were incubated with primary antibody GluA2 (AGC-005, Alomone Labs) was for 2 hours. Protein A coupled to 10nm colloidal gold particles was used to target the primary antibody (G. Posthuma, University Medical Center Utrecht) (1:50).

### Patch-clamp recordings from EVs-treated hippocampal neurons

Whole-cell patch clamp measurements^37^ were performed on DIV12-14 hippocampal neurons. Neuron cultures were prepared from mouse embryos (E.16) as described above. Pipettes were made from borosilicate glass and had resistances of 3 - 5 MΩ after filling with intracellular solution (120 mM K-gluconate, 8 mM NaCl, 2 mM MgCl_2_, 0.5 mM CaCl_2_, 5 mM EGTA, 10 mM HEPES, 14 mM phosphocreatine, 2 mM magnesium-ATP, 0.3 mM sodium-GTP, and pH adjusted to 7.3 with KOH). Patchmaster software (HEKA, Lambrecht, Germany) and an EPC-9 patch-clamp amplifier (HEKA) were used for data acquisition and pulse application. Recordings were low-pass filtered at 2.9 kHz and analyzed with Fitmaster (HEKA, Lambrecht, Germany), Igor Pro 6.03 (Wavemetrics), and Excel (Microsoft). Neurons with an access resistance <20 MΩ were evaluated. Current clamp recordings were performed on untreated cultured neurons and on neurons exposed to extracellular vesicles. Current clamp experiments were done at room temperature (21-23°C) in Ringeŕs solution (143 mM NaCl, 5 mM 1 KCl, 0.8 mM MgCl_2_, 1 mM CaCl_2_, 10 mM HEPES, 5 mM glucose, and pH adjusted to 7.3 with NaOH). All substances were purchased from Sigma Aldrich.

### Mass spectrometry-based bottom-up proteomics

#### Tryptic Digestion

Samples were dissolved in 100 mM triethyl ammonium bicarbonate (TEAB) and 1% w/v sodium deoxycholate (SDC) buffer, boiled at 95 °C for 5 min and sonicated with a probe sonicator. The protein concentration of denatured proteins was determined by the Pierce bicinchoninic acid assay (BCA) Protein assay kit (Thermo Fisher) and samples were diluted to 10 µg of protein in 25 µL buffer. The samples were then pipetted into a 96-well LoBind plate (Eppendorf, Hamburg, Germany) placed on an Andrew+ Pipetting Robot (Waters, Milford, USA), which was used to execute all following steps. Using the robot, disulfide bonds were first reduced in 10 mM dithiothreitol for 30 min at 56 °C while shaking at 800 rpm and alkylated in presence of 20 mM iodoacetamide for 30 min at 37 °C, again while shaking at 800 rpm. Then, carboxylate modified magnetic E3 and E7 speed beads (Cytvia Sera-Mag™, Marlborough, USA) at 1:1 ratio in LC-MS grade water were added in a 10:1 (beads/protein) ratio to each sample, following the single-pot, solid-phase enhanced sample preparation (SP3)-protocol workflow 1. To bind the proteins to the beads, acetonitrile (ACN) concentration was raised to 50%. Subsequently, samples were shaken at 600 rpm for 18 min at room temperature. Magnetic beads were magnetized, and the supernatant was removed. Magnetic beads were further washed two times with 80% Ethanol (EtOH) and then two times with 100% ACN. After resuspension in 100 mM AmBiCa, digestion with trypsin was performed (sequencing grade, Promega) at 1:100 (enzyme:protein) ratio at 37 °C overnight while shaking at 500 rpm. The next day, trifluoroacetic acid (TFA) was added to a final concentration of 1% to inactivate trypsin. The samples were then shaken at 500 rpm for 5 min at room temperature. Finally, beads were magnetized, and the supernatant containing tryptic peptides was transferred into a new 96-well LoBind plate, ready for subsequent LC-MS/MS analysis.

### LC-MS/MS measurement

Chromatographic separation of tryptic peptides was achieved with a two-buffer system (buffer A: 0.1% FA in H2O, buffer B: 0.1% FA (Fisher Chemical, A117-50) in 80% ACN) on a UHPLC (VanquishTM neo UHPLC system, Thermo Fisher). Attached to the UHPLC was a peptide trap (300 µm x 5 mm, C18, PepMap™ Neo Trap Cartridge, Thermo Fisher, 174500) for online desalting and purification, followed by a 25 cm C18 reversed-phase column (75 µm x 250 mm, 130 Å pore size, 1.7 µm particle size, peptide BEH C18, nanoEase, Waters, 186008795). Peptides were separated using an 80 min method with linearly increasing ACN concentration from 2 % to 30 % ACN over 60 minutes. MS/MS measurements were performed on a quadrupole-orbitrap hybrid mass spectrometer (Exploris 480, Thermo Fisher Scientific). Eluting peptides were ionized using a nano-electrospray ionization source (nano-ESI) with a spray voltage of 1,800 and analyzed in data independent acquisition (DIA) mode. For each MS1 scan, ions were accumulated for a maximum of 240 milliseconds or until a charge density of 3 x 10 6 ions (AGC Target) was reached. Fourier-transformation based mass analysis of the data from the orbitrap mass analyzer was performed covering a mass range of m/z 400 – 1,400 with a resolution of 120,000 at m/z 200. Within a precursor mass range of m/z 380-980 fragmentation in DIA-mode with m/z 12 isolation windows and m/z 1 window overlaps was performed. Fragmentation was performed at normalized collision energy of 28% using higher energy collisional dissociation (HCD). An AGC target of 2 x 10 6 ions or a maximum of 54 ms was set. Orbitrap resolution was set to 30 000 with a scan range from m/z 350-2000.

### LC-MS/MS data processing and analysis

LC-MS/MS data were searched with the CHIMERYS DIA algorithm integrated into the Proteome Discoverer software (v3.1.0.638, Thermo Fisher Scientific) against a reviewed murine Swissprot database using Inferys 3.0 fragmentation as prediction model. Carbamidomethylation was set as a fixed modification for cysteine residues. The oxidation of methionine was allowed as variable modification. A maximum number of one missing tryptic cleavage was set. Peptides between 7 and 30 amino acids were considered. A strict cutoff (FDR < 0.01) was set for peptide identification. Quantification was performed by CHIMERYS based on fragment ions.

For the statistical and bioinformatics analyses of the proteomics data sets, we relied on base R and various packages in R. Obtained protein abundances were log 2 -transformed and normalized by column-median. Proteins which were detected in least four replicates of a group (EV preparations and cell lysates individually) were kept for further analysis. Proteins were considered to be differentially regulated if the Student’s *t*-test-based *p*-value was ≤ 0.05 and at least a 1.5-fold change in either direction was observed. Proteins were considered uniquely present in a certain group when they were expressed in at least four out of six replicates for that particular group and were simultaneously absent in the other group during pairwise comparison.

Venn diagram was prepared using *BioVenn* package (version 1.1.3, 2021). Correlation plots were prepared using *ggcorrplot* package (version 0.1.4, 2021). Pearson’s coefficient values were obtained using base R. Initially, proteins with missing values were excluded from the Principal component analyses and heatmaps. Principal component analysis (PCA) was carried out using the *prcomp* function of the base R. Resultant PCA plot was composed utilizing the *ggplot2* package (version 3.3.3, 2016). Volcano plot was composed using *EnhancedVolcano* package (version 1.16.0, 2021) in R Studio.

Gene ontology (GO) analyses were performed utilizing the *clusterProfiler* (version 4.8.1, 2024), *enrichR* (version 3.2, 2024), *org.Mm.eg.db* (version 3.16.0, 2023), and *topGO* (version 2.50.0, 2023) packages. Overrepresentation analysis (ORA) was performed to find associations of differentially upregulated proteins with the GO category ‘Biological Process’ with the following analysis parameters: *p*valueCutoff = 0.05, *q*valueCutoff = 0.2, *p*AdjustMethod = “BH”, minGSSize = 5, and maxGSSize = 500. Reactome pathway enrichment analyses were conducted using *ReactomePA* package (version 1.14.0, 2021), using ‘mouse’ database as reference with following analysis parameters: *p*valueCutoff = 0.05, *p*AdjustMethod = “BH”, *q*valueCutoff = 0.2, minGSSize = 10, and maxGSSize = 50. Protein interaction network was prepared using string-db.org.

## Acknowledgements

We thank Christoph Krisp, Katharina Kolbe, Chudamani Raithore and Yvonne Pechmann for their technical support.

## Author contributions

F.L.L., M.S., A.M-A., M.K. and M.G. planned the experiments. F.L.L., J.R.S., M.S., A.M-A., D.S., K.V.G., C.K., L.B., M.S., performed the experiments and analyzed the data. F.L.L., A.M-A., M.S., and M.G. wrote the manuscript. F.L.L., J.R.S., D.S., K.V.G., C.K., M.S., A.M-A., L.B., M.S., T.M., M.G., M.K., F.L.R., H.S. and A.F.H. reviewed and edited the manuscript.

---

**Supplementary Figure S1:**
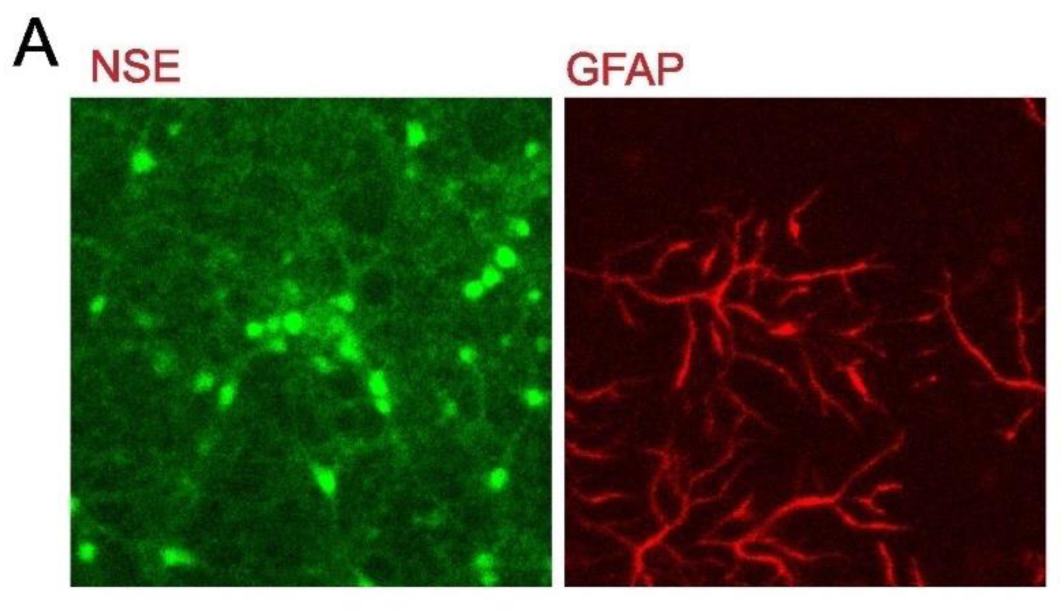
Primary cultures from the mouse cortex contain both neurons and astrocytes. A) Immunocytochemistry against Neuronal Specific Enolase (NSE) and GFAP (astrocytes) depicts the presence of both cell types.

**Supplementary Figure S2:**
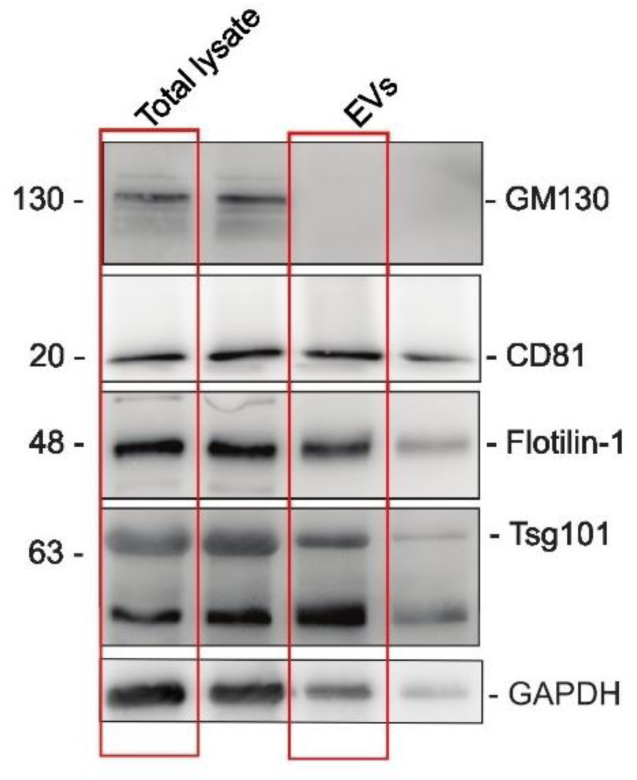
Related to Fig. 2C. Westernblot showing the lanes selected for visualization in Fig. 2C.

**Supplementary Figure S3:**
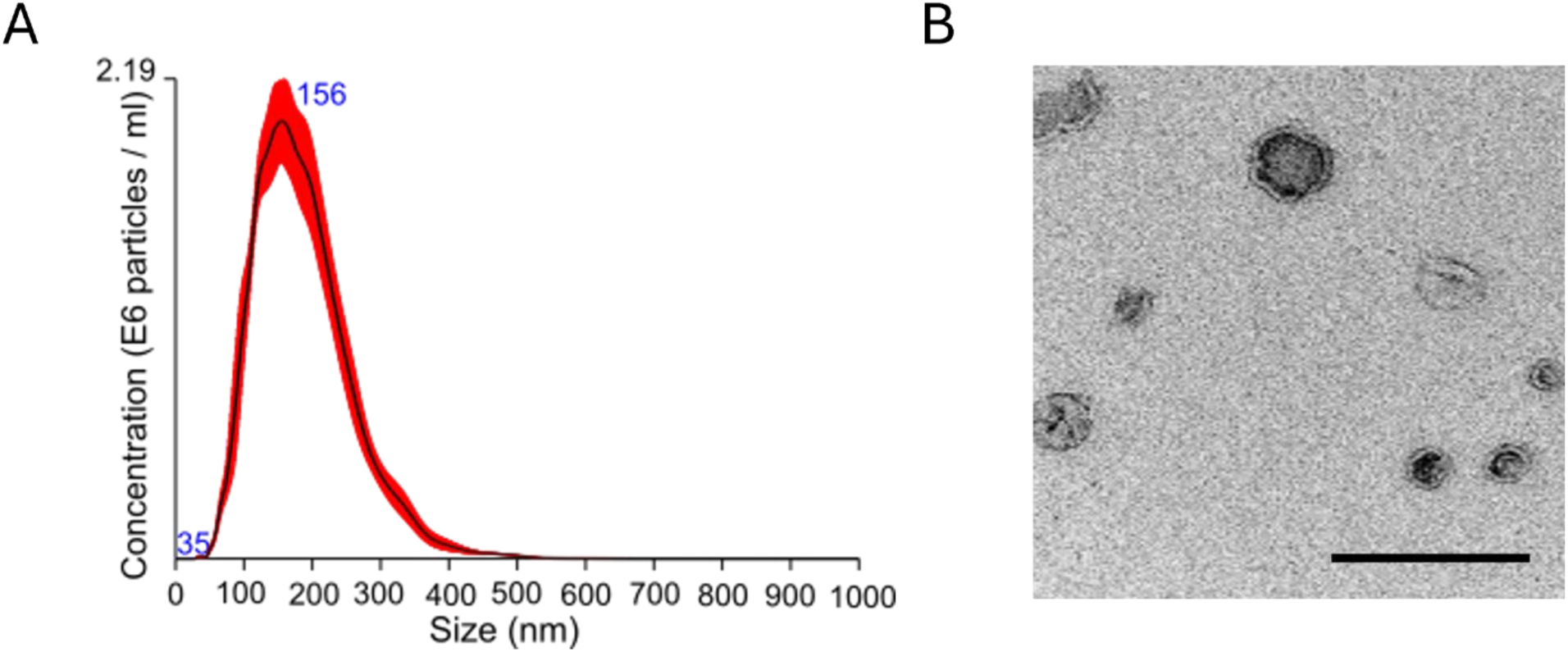
The BDEVs show the expected size distribution profile and morphology: A) Representative NTA of the BDEVs shown in Figure 4. The BDEVs showed the usual normal-like size distribution with an average diameter of 155 nm. B) Representative TEM image of the BDEVs showing the membrane of the vesicles and their characteristic cup-shape. Scale bar = 200nm.

